# ECLiPSE: A Versatile Classification Technique for Structural and Morphological Analysis of Super-Resolution Microscopy Data

**DOI:** 10.1101/2023.05.10.540077

**Authors:** Siewert Hugelier, Hannah Kim, Melina Theoni Gyparaki, Charles Bond, Qing Tang, Adriana Naomi Santiago-Ruiz, Sílvia Porta, Melike Lakadamyali

## Abstract

We introduce a new automated machine learning analysis pipeline to precisely classify cellular structures captured through single molecule localization microscopy, which we call ECLiPSE (Enhanced Classification of Localized Pointclouds by Shape Extraction). ECLiPSE leverages 67 comprehensive shape descriptors encompassing geometric, boundary, skeleton and other properties, the majority of which are directly extracted from the localizations to accurately characterize individual structures. We validate ECLiPSE through unsupervised and supervised classification on a dataset featuring five distinct cellular structures, achieving exceptionally high classification accuracies nearing 100%. Moreover, we demonstrate the versatility of our approach by applying it to two novel biological applications: quantifying the clearance of tau protein aggregates, a critical marker for neurodegenerative diseases, and differentiating between two distinct morphological features (morphotypes) of TAR DNA-binding protein 43 proteinopathy, potentially associated to different TDP-43 strains, each exhibiting unique seeding and spreading properties. We anticipate that this versatile approach will significantly enhance the way we study cellular structures across various biological contexts, elucidating their roles in disease development and progression.

## Main

Cells are compartmentalized into various structural units including membrane bound and membraneless sub-cellular organelles, cytoskeletal structures, and supramolecular protein assemblies. Each one of these organizational units possesses unique and complex structural and morphological properties that span a range of length scales to match their function. The distinct morphology of organelles help adopt them to specific functions, and can change in response to cellular needs as well as in disease states^1^. Similarly, aggregation of proteins into solid inclusions with specific morphological properties is a hallmark of several neurodegenerative diseases^2^. Therefore, techniques to characterize and classify subcellular compartments based on their structural and morphological properties are invaluable in studying both cell physiology and pathology.

Recent advancements in super-resolution microscopy have revolutionized our ability to visualize the intricate morphological features of sub-cellular compartments and organelles at nanoscale spatial resolution^3^. Super-resolution microscopy can capture subtle changes in the morphology and structure of these subcellular components, which were once inaccessible due to the diffraction limit of light. For this purpose, Single Molecule Localization Microscopy (SMLM) techniques have been widely adopted by the cell biology community, as they do not require highly specialized microscope hardware^4^. However, the development of analysis tools to accurately classify individual sub-cellular structures into distinct categories based on shape and morphology has not kept pace with advancements in super-resolution microscopy, particularly in the context of SMLM. This is because SMLM data consist of point clouds rather than pixelated intensity-based images, which is less compatible with traditional image processing and analysis techniques.

Current tools employ template-based or template-free strategies to classify and align super-resolution data of highly symmetric, self-similar, and frequently simplistic structures, such as the nuclear pore complex (NPC), to facilitate single particle averaging for the purpose of accurately describing the structure of interest ^5-9^. However, these methods do not explicitly ascertain the morphological characteristics of unique structures. Alternative tools, such as LocMoFit, depend on model fitting with pre-defined and often basic geometric models to achieve the structure-matching and extract basic quantitative properties^10^. LocMoFit enables the extraction of a number of geometric features that are included in the model, such as size and symmetry angle, to determine the degree of variability among individual structures. Although this method is valuable, it can only quantify a small number of parameters and cannot been utilized to classify structures into distinct categories, as it relies on analyzing structures that are identical or highly similar. Recently, automated structure analysis program (ASAP) was developed to quantify and classify structures based on a limited number of geometric shape descriptors^11^. ASAP was applied to SMLM images of NPCs, endocytic vesicles and Bax protein pores, all of which assemble into small (∼100 nm) and simple structures resembling either rods, arcs or circles. Although ASAP represents a significant improvement in the structural classification of super-resolution data, one of its drawbacks is that it requires rendering the SMLM point cloud data into pixelated images, which may introduce artefacts and cause the loss of important information. Furthermore, the limited number of shape descriptors in ASAP makes it less applicable to structures with complex shapes and larger sizes.

To fill the gaps in advanced classification tools, we developed an analysis pipeline called ECLiPSE (Enhanced Classification of Localized Pointclouds by Shape Extraction) that expands the toolbox for classifying structures in SMLM data (Figure 1). ECLiPSE can accurately describe and classify structures that range in size between < 100 nm to several microns and possess a high degree of structural complexity, including organelles with distinct morphologies, cytoskeletal filaments, and diverse protein aggregates. After data acquisition and segmentation of super-resolution data (which can be done using existing techniques and packages, e.g., Voronoi tessellation, or DBScan^12-15^;Figure 1A, B), the first step in ECLiPSE involves calculating 67 shape descriptors, of which the majority is extracted directly from the point cloud data (Figure 1C). The shape descriptors include geometric properties, boundary properties, skeleton properties, texture properties, Hu moments, and fractal properties (Supplementary Table 1). It is important to note that not every shape descriptor is equally effective at distinguishing between various structures and the specific descriptors that provide the highest degree of separation can vary depending on each biological application. To address this, we have incorporated an automated variable selection step (Supplementary Note 1 and Supplementary Figure 1), which uses *a priori* information on the classes to select the most informative descriptors that distinguish between different classes (Figure 1D). If *a priori* information on the classes is not available, data compression algorithms such as Principal Component Analysis (PCA) could be used to extract such information. Once the shape descriptors are calculated and optionally undergo variable selection, the data can be explored in the PCA-space using these quantitative features (Figure 1E). This approach offers a preliminary visual representation of the extent to which datasets are separated within the PCA space and can uncover sub-populations within the data. Additionally, *a priori* information about distinct groups can be employed to color-code them, revealing their degree of separation. The final step is the classification (Figure 1F) using many machine learning models including supervised models (e.g., K-nearest neighbors, Random Forest, Partial Least Squares for discrimination, Logistic Regression – Discriminant Analysis, etc.) and unsupervised models (e.g., partial or agglomerative hierarchical clustering). Model validation is performed on a subset of training data, or ground truth data that was not incorporated during the initial training phase. The training and validation process can be performed multiple times and the best performing model(s) is (are) then automatically selected to predict the class membership of new data that the user provides. More information on the unsupervised and supervised classification and hyperparameters used in this work can be found in Supplementary Note 2, Supplementary Figures 2, 3, and Supplementary Table 2.

**Figure 1:**
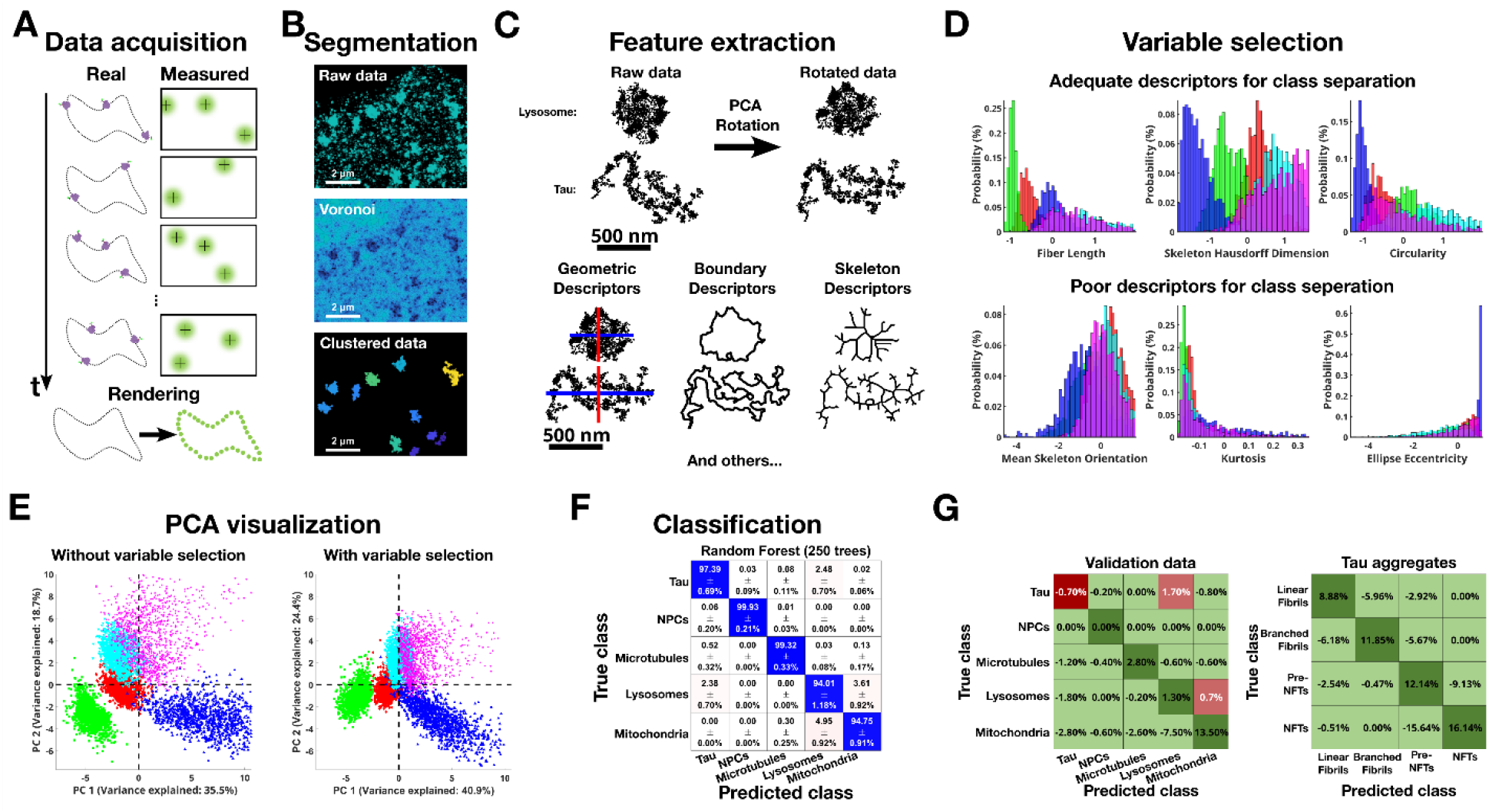
The analysis pipeline of ECLiPSE to quantify super-resolution microscopy data as point-clouds applied to the validation data. (A) Schematic representation of the SMLM data acquisition process; (B) Segmentation of the localizations into individual clusters (applied to an ROI of the lysosome data); (C) Feature extraction from segmented point-cloud clusters generate geometric, boundary, skeleton, etc. descriptors; (D) Example distributions of features that adequately or poorly separate the different classes in the validation data, as determined by automatic variable selection (28/67 descriptors that provide clear class separation; Red: Tau, Green: NPCs, Blue: Microtubules, Cyan: Lysosome, Magenta: Mitochondria); (E) Data exploration using PCA, with and without variable selection (Red: Tau, Green: NUP96, Blue: Microtubules, Cyan: Lysosome, Magenta: Mitochondria); (F) Optimized classification results for the validation data (97.1 ± 0.1% accuracy), obtained by the Random Forest classifier (100 best models out of 1000 generated models); (G) Difference confusion matrices between ECLiPSE (Logistic Regression, no variable selection) and ASAP (10nm rendering precision, 1.5 ×10^5^ threshold, Discriminant classifier). Left panel: validation data (96.9% vs 93.5% accuracy for ECLiPSE and ASAP, respectively), right panel: Tau aggregation data (92.9% vs 80.6% accuracy for ECLiPSE and ASAP, respectively). Green values represent superior results for ECLiPSE (i.e., positive diagonal values and negative off-diagonal values), whereas red values represent inferior results for ECLiPSE (i.e., negative diagonal values and positive off-diagonal values).

We first validated our approach using ground truth SMLM datasets of five distinct structures including organelles (lysosomes, mitochondria), cytoskeletal filaments (microtubules), supramolecular assemblies (nuclear pore complex)^16^ and aggregates of the tau protein (Supplementary Materials and Methods, and Supplementary Figure 4). Performing variable selection on this dataset largely reduced the variance between members of the same class, as shown by the exploratory PCA analysis but did not significantly improve class separation (Figure 1E and Supplementary Video 1). We then trained several types of machine learning models using a limited subset of training data (approximately 550 samples per group) and subsequently made predictions using ground truth data that had been excluded from the training dataset. The best results were achieved with the Random Forest classifier with an average prediction accuracy of 97.1 ± 0.1% across all categories (Figure 1F; the prediction accuracy is the average percentage of correctly classified data over all classes).

The largest confusion in the prediction was between lysosomes and mitochondria (94.0 ± 1.2% and 94.8 ± 0.9%, respectively) as these classes are morphologically more similar to each other than to the other classes. We thus selected these two classes for a side-by-side comparison of our approach to the previously developed ASAP (Supplemental Note 3, Supplementary Figures 5, 6, and Supplementary Table 3). Since ASAP does not provide an automatic model selection, we used the classification method included as default setting (Discriminant Analysis). Moreover, given that ASAP requires pixelated images, we also tested how its performance depends on the image rendering parameters, in particular the width of the rendering Point Spread Function (PSF) and binary image threshold (Supplementary Figure 5). The accuracy of the ASAP prediction was dependent on both parameters as expected (Supplementary Figure 6), and these parameters must therefore be manually optimized to achieve maximal results. Additionally, a full study on the influence of these parameters on ASAP classification accuracy for all available methods was performed. Surprisingly, it revealed that the relationship between rendering PSF width and prediction accuracy was model dependent. Sometimes, a larger rendering PSF size led to more accurate predictions, but for other classification methods, smaller PSF sizes resulted in superior performance (Supplementary Table 3). Moreover, upon comparing ASAP to ECLiPSE using their default settings, ECLiPSE demonstrated a superior average prediction accuracy by 6.1% over ASAP, and ECLiPSE also performed better than the best average prediction accuracy achieved in the optimized study presented in Supplementary Table 3. A similar result is also obtained when utilizing the validation dataset including all five classes (Figure 1G, left), where the difference in average prediction accuracy is 3.4%, with a maximum difference in prediction accuracy of 13.5% for the more heterogeneous mitochondria class. Furthermore, with ASAP, a significant disparity in prediction accuracy was observed across various classification methods, whereas this inconsistency was not present when employing ECLiPSE (Supplementary Figure 3). These results demonstrate several advantages of our approach over existing tools: the ability to use the unbiased raw point cloud data, automated variable selection and automated model selection. These collectively provide improved performance and robustness over previous state-of-the-art methods.

We next applied our approach to two biological applications: clearance of tau protein aggregates (Figure 2A-E) and detection of TAR DNA-binding protein 43 (TDP-43) proteinopathy morphotypes (Figure 2F-I). Both applications represent aggregation of proteins into insoluble inclusions that play a role in several neurodegenerative diseases^17^. Tau is a neuronal microtubule associated protein, which undergoes aberrant post-translational modifications and aggregation in several tauopathies including frontotemporal dementia with Parkinsonism linked to Chromosome 17 (FTDP-17), Pick’s Disease and Alzheimer’s Disease (AD)^18^. It has been shown that these tau inclusions are morphologically diverse and disease-specific (for example neurofibrillary tangles or NFTs in AD and Pick’s bodies in Pick’s Disease)^19^. Recent work suggests that there are molecularly and structurally distinct disease specific tau strains in which the tau protofilaments assume a distinct fold that leads to disease specific tau aggregation^19-22^. However, the relationship between the molecular signatures (e.g., post-translational modifications) of tau proteins, the tau protofilament structure and the morphology of the resulting tau aggregates is not clearly understood. Using SMLM, we previously showed that tau forms morphologically diverse aggregates in an FTDP-17 engineered cell model^23^. These aggregates were broadly categorized into four classes based on visual inspection: linear fibrils, branched fibrils, pre-neurofibrillary tangles (pre-NFTs) and NFTs^23^ (Figure 2A). Interestingly, these aggregate classes were enriched with hyper-phosphorylation marks on distinct tau residues (phospho-Ser202/205 for linear fibrils and NFTs; and phospho-Thr231 for branched fibrils)^23^. The automated, high-throughput, and unbiased classification of these previously identified tau aggregates is crucial for obtaining insights into the progression of tau pathology. However, this task is particularly challenging due to the irregular and highly diverse morphological features of these aggregates. We applied our machine learning classification and validated that ECLiPSE accurately discriminates between members of the four different tau morphological classes (Figure 2B, C). The average prediction accuracy was 89.8 ± 0.4%, which is remarkably high given the high complexity and morphological similarity among the different tau aggregate structures. Additionally, we compared ECLiPSE and ASAP on this complex dataset and found that, at default settings for both methods, ECLiPSE yielded a 12.3% increase in overall prediction power relative to ASAP when accounting for all aggregate classes. (Figure 1G, right, and Supplementary Table 4). Notably, ECLiPSE demonstrated an impressive 17.7% improvement in prediction accuracy for some of the most challenging comparisons (Supplementary Table 4). Once again, these results underscore the robustness of our approach in handling challenging biological data where the morphology of structures is complex and spans a broad size scale. To determine the contribution of the automated variable selection to the high performance, we repeated the prediction without variable selection and found that this step was indeed important to improve both prediction accuracy and its robustness (Supplementary Figure 7).

**Figure 2.**
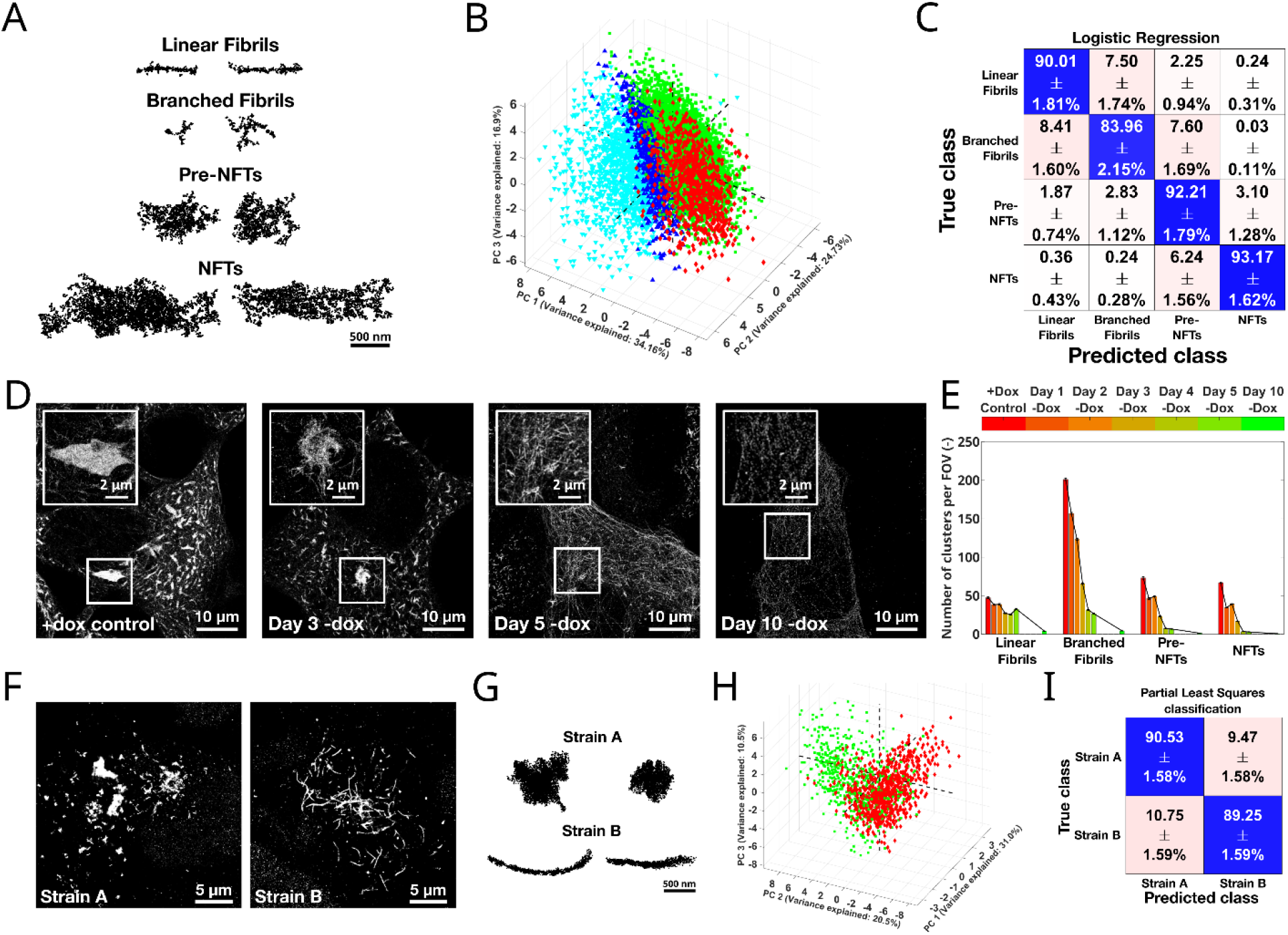
Biological applications with ECLiPSE. (A) Example clusters of the four different tau aggregate species (i.e., Linear Fibrils, Branched Fibrils, pre-NFTs, and NFTs); (B) Data exploration using PCA indicates that the data are complex and classes cannot easily be separated (Red: Linear Fibrils, Green: Branched Fibrils, Blue: pre-NFTs, Cyan: NFTs); (C) Classification results obtained (89.8 ± 0.4% accuracy) using the Logistic Regression classifier on the variable selected tau aggregates data (100 best models out of 1000 generated models); (D) Tau clearance, after removal of doxycycline, observed over a period of 10 days shows a visual reduction of tau aggregate cluster sizes over time; (E) Tau aggregate species prediction on the total dataset demonstrates the rapid decrease in Branched Fibrils, pre-NFTs and NFTs starting at day 1 after doxycycline removal, whereas Linear Fibrils show delayed degradation kinetics; (F) Acquired SMLM data of two patient specific TDP-43 strains; (G) Representative clusters of the two patient specific TDP-43 strains; (H) Data exploration using PCA indicates that a non-negligible subset of the data clusters are similar between the two patient specific strains (Red: Strain A; Green: Strain B); (I) Classification results obtained (89.9 ± 0.6% accuracy) using the Partial Least Squares classifier on the non-variable selected TDP-43 data (100 best models out of 1000 generated models). For (E) +Dox control: n = 29 cells, Day 1 -Dox: n = 27 cells, Day 2 -Dox: n= 30 cells, Day 3 -Dox: n = 27 cells, Day 4 -Dox: n = 29 cells, Day 5 -Dox: n = 27 cells, Day 10 -Dox: n = 28 cells. Error bars represent the standard deviation from the prediction of the 100 best models as shown in (C).

Following this validation, we next examined data in which we induced tau degradation. To do so, we inhibited the expression of soluble tau by removing doxycycline (Dox) in the QBI-293 (Clone 4.1) cells (Figure 2D and Supplementary Materials and Methods). We observed a drastic decrease in total tau amounts and tau aggregates at days 1 through 10 after removing Dox (Figure 2D and Supplementary Figure 8), which is consistent with previous biochemical analysis^24^. Previous work had shown that this loss corresponds to tau degradation mediated by both the proteasome and autophagy pathways, but it remains unclear how the different tau aggregate classes are cleared over time. Using ECLiPSE, we predicted the number of the four morphological tau aggregate classes at different time points following Dox removal, which allowed us to determine the timing of degradation of these different classes (Figure 2E). Interestingly, we found that while branched fibrils, pre-NFTs and NFTs showed a rapid and consistent decay starting at day 1 after Dox removal, linear fibrils persisted up to day 5 displaying a more delayed degradation kinetics (Figure 2E). These results suggest that it may be more challenging to clear linear fibrils compared to other morphological tau aggregate classes. Alternatively, it is possible that other aggregate classes are broken down into linear fibrils resulting in their accumulation over time. Since both autophagy and proteasome pathways are involved in tau aggregate clearance^24^, in the future, this approach would be useful to determine if specific pathways clear distinct classes of tau aggregates and the mechanisms of why the linear fibrils have a delayed degradation kinetics.

Finally, we used our approach to discriminate between brain-derived TDP-43 strains, obtained from two patients with frontotemporal lobular degeneration with TDP-43 immunoreactive pathology (FTLD-TDP). In normal conditions, TDP-43 is found in the nucleus and plays an important role in RNA regulation^25^. In pathology, changes in cleavage and post-translational modifications of TDP-43 lead to its cytoplasmic accumulation and aggregation into inclusions, similar to tau^25^. Previous work has demonstrated that extracts derived from the postmortem brain samples of individuals with FTLD-TDP can seed morphologically distinct TDP-43 aggregates in both animal and cell models^26^. This finding supports the existence of distinct TDP-43 strains that possess unique seeding and spreading properties, which is highly relevant to understanding the pathophysiology of the disease. However, previous work has relied on low resolution images and simple geometric measurements (e.g., circularity) to distinguish between “globular-like” versus “wisp-like” TDP-43 aggregates seeded by these different strains. Manual measurements and classification of low-resolution images can be a slow and subjective process. While this approach is useful, super-resolution information is needed to precisely visualize and quantify the morphology of TDP-43 aggregates and robustly classify them. We thus aimed to apply ECLiPSE to determine if this approach could detect morphologically distinct TDP-43 aggregates in cell models. TDP-43 extracted from two distinct postmortem FTLD-TDP brains was used to seed TDP-43 aggregates in cell models. The resulting aggregates were acquired using SMLM (Figure 2F, G and Supplementary Materials and Methods). Visual inspection confirmed that one strain led to the formation of more globular-like aggregates (Figure 2F, G – Strain A), whereas the other strain predominantly seeded aggregated that resembled linear fibrils, previously described as wisps (Figure 2F, G – Strain B). Extracting shape descriptors enabled us to further confirm these differences in morphology using PCA analysis (Figure 2H). Finally, we applied the machine learning classification on the clustered localization data of aggregates from the two TDP-43 strains not included in the training data and showed that ECLiPSE predicts the distinct morphologies with very high accuracy (89.9 ± 0.6%) (Figure 2I). Furthermore, upon examining the clusters that were accurately or inaccurately classified (Supplementary Figure 9), it became evident that the “misidentified” clusters of one strain exhibited morphological features characteristic of the other strain, and vice versa. We can therefore conclude that although a strain primarily seeds aggregates with a specific morphological trait, a considerable proportion of the seeded aggregates still exhibits morphological similarities to the other strain even when the strains are derived from two distinct postmortem FTLD-TDP brains.

These results demonstrate the ability of ECLiPSE to classify the presence of morphologically distinct protein aggregates in postmortem brain tissue and link aggregate morphology to patient-specific proteinopathy strains. Importantly, ECLiPSE is not limited to classifying the morphology of protein aggregates; we envision that it will also be broadly applicable for classifying changes in organelle morphology, cytoskeletal architecture, and other supramolecular assemblies in the context of the cell. Additional applications of ECLiPSE include assigning pseudo-time stamps to protein aggregates or other biological structures based on their evolving morphological properties, as well as generating pseudo-multi-color super-resolution images by color-coding distinct structures within single-color images. Overall, we have developed a toolbox with broad applicability in structural and morphological characterization of super-resolution microscopy data.

## Data availability

An example dataset of the validation data on which the ECLiPSE can be tested is provided with the source code. The remainder of the data that supports the findings of this work are available from the corresponding authors upon request.

## Code availability

The source code, example scripts and documentation of ECLiPSE can be found on https://github.com/LakGroup/ECLiPSE.

## Acknowledgements

We thank S. Thakur for providing super-resolution data of the nuclear pore complex. M.L. acknowledges funding from NIH RO1 GM133842, RM1 GM136511, UO1 DA052715 and RO1 AR079224.

## Author contributions

S.H. and M.L. conceived and designed the research study, S.H. and H.K. designed the ECLiPSE algorithms and pipeline, M.T.G., C.B., Q.T., A.N.S.-R. and S.P. performed experimental work and provided data, S.H. performed the analysis with assistance from H.K., S.H. assessed method performance, S.H. and M.L. wrote the manuscript with input from all other authors.

