## Supplementary Information for "ECLiPSE: A Versatile Classification Technique for Structural and Morphological Analysis of Super-Resolution Microscopy Data"

### Supplementary Note 1 – Variable selection methods

#### General information

Variable selection methods provide ways to determine how relevant and/or informative single variables (shape descriptors in this work, see Supplementary Table 1) are and have as a goal to maximize model performance, minimize noise introduction by overfitting, remove correlation and make analysis more robust. They are an important field of research in spectroscopy (for both prediction and classification), where identification of the relevant variables also provides empirical knowledge about the process under investigation[1-3]. Typically, most of these methods are based on adding or removing variables from the model and evaluating their influence on its performance. Practically speaking for variable selection in classification, it is also possible that the addition or removal of a variable is to focus on the separation of a single class from the others, rather than all classes together, but this will eventually lead to better model performance. An observation of its influence can for example be seen when exploring the data with Principal Component Analysis (PCA): e.g., the variable selected data explains more data variance using fewer principal components, or the within-class variance is reduced (i.e., more 'compact' in the PCA space). Furthermore, its influence is manifested during classification as an improvement in average classification accuracy, and/or the standard deviation in the classification confusion matrices is decreased with respect to using the full data set (i.e., improved robustness of the classification). For a more complete information on the benefits of variable selection, we refer to references[1,2].

It is worth noting that several of the used classification methods in this work include an indirect variable selection. For example, any classification method that uses PCA compression before classifying the data will have a (limited) variable selection. Examples of these routines are Partial Least Squares classification or Logistic Regression (can be both with and without PCA or Partial Least Squares, PLS, compression). Random Forest classification on the other hand does not use PCA to compress the data, but it randomly selects a subset of the variables to construct its decision trees which can be used to investigate their prediction power in the global model, and it is therefore also capable of providing information on which variables are important to correctly classify the data.

Since not every classification method inherently incorporates variable selection capabilities, we have deliberately integrated this feature into the Enhanced Classification of Localized Pointclouds by Shape Extraction (ECLiPSE) analysis pipeline. Moreover, given that interpreting significant variables within the PCA/PLS space can be complex and unintuitive, this deliberate incorporation of variable selection empowers researchers to examine the influence of diverse variables more effectively in discerning distinct structural classes.

#### Implementation and results

This work includes ten different variable selection methods, of which six methods (biPLS – Backwards interval PLS[4], rPLS – Recursive PLS[5], Chi Square test[6], MRMR – Minimum Redundancy Maximum Relevance[7], ReliefF[8], and Boruta[9]) are available as either a built-in function of the MATLAB Statistics & Machine Learning Toolbox, or provided as stand-alone functions in the code. The remaining four methods (biPLS – Backwards Interval PLS[4], rPLS – Recursive PLS[5], iPLS – Interval PLS[4] & GA – Genetic Algorithm[10]) are included in the PLS Toolbox (Eigenvector Inc., free trials are available on their website for academics). Two of the methods have multiple implementations (biPLS and rPLS) and may lead to somewhat different results. This is related to slight differences in their implementation and selection criteria. In our experience, the selection criteria for the PLS toolbox routines are stricter, as shown in Supplementary Figure 1A. Nonetheless, it is important to highlight that all variables chosen by the PLS Toolbox implementation are also selected in the stand-alone implementations of the corresponding methods, along with additional variables.

Variable selection within ECLiPSE is performed by applying it to a subset of the data and repeating this process multiple times (in this work, it was repeated 250 times for all data sets). Using this method, frequency plots of how often a variable was selected by a given method are obtained (such as shown in Supplementary Figure 1A). This provides the necessary information to do the variable selection with a higher robustness to outliers in the data. This implementation is especially useful when the within-class variance is large, which is the case in the data presented in this work as none of the structures, besides the Nuclear Pore Complex structures, are highly symmetrical and uniform. A threshold is used to determine the cutoff of useful variables (Supplementary Figure 1A, B; indicated by the dashed line).

Ten different variable selection methods are available in ECLiPSE, and their results were combined to further increase the robustness of the variables selected (see Supplementary Figure 1B). Practically speaking, the number of methods used for the validation data is 8 (the Chi Square and GA method did not provide useful results), for the tau aggregates also 8 (the MRMR and Chi Square methods did not provide useful results), and for the TDP-43 data all 10 methods were used. A threshold was set accordingly, as indicated by the dashed line in each panel. The total number of important variables for each data set is 28, 27 and 22, respectively.

### Supplementary Note 2 – Classification methods

#### General information

Classification methods can broadly be categorized into two groups: unsupervised classification, or clustering, and supervised classification. Unsupervised classification aims at grouping similar samples without relying on any prior knowledge of class memberships within the data, while supervised classification leverages known class information to construct its models. Unsupervised classification is advantageous because it identifies the natural clusters inherent in the data, providing insights into the similarity among various samples. However, since it does not utilize any prior information to optimize discrimination among different groups in the data, it often leads to inferior models compared to supervised approaches. Consequently, unsupervised classification is commonly employed as an exploratory analysis tool rather than for definitive classification purposes.

In the main text of this work, supervised classification approaches are presented, because their prediction power is higher than for the unsupervised methods, but ECLiPSE is not limited to using supervised classification. Oftentimes, classes are not known beforehand, or the clustering approaches can be used to detect and uncover natural grouping of the data (e.g., subgroups within a group).

#### Unsupervised classification

In this work, unsupervised classification algorithms based on Hierarchical Clustering Analysis (HCA)[11] are included in the available methods, by using implementations of the PLS toolbox (Eigenvector Inc.). Built-in MATLAB methods available in the Statistics and Machine Learning Toolbox (e.g., k-means[12]) can be used as well. However, the advantage of using the PLS toolbox for this purpose is that it allows to easily visualize the results and inspect the clustering process.

An extensive study on the different available HCA methods of the PLS Toolbox was performed, on all the different data sets showcased in this work. Partitional clustering (i.e., methods that progressively divide clusters into smaller ones) provided less accurate results than agglomerative clustering methods (i.e., methods that progressively combine clusters into bigger ones), and will therefore not be presented. Within the scope of the work presented here, different distances to link the different clusters together were explored (nearest neighbor, furthest neighbor, pair-group average, centroid, median, and Ward's method), but only the results obtained by the Ward's method for linkage are shown as they were superior to other distances explored. Other parameters that were optimized were: variable selection (yes/no) – PCA transformation (yes/no) – number of principal components (if performing the clustering on PCA-transformed data, with the maximum being the number of variables in the data) – Distance metric (Euclidean/Mahalanobis when using PCA transformed data, Manhattan/Euclidean distance when not using PCA transformed data).

The best results that emerged from this optimization study are presented in Supplementary Figure 2. Although the classification accuracy is lower compared to those achieved by supervised approaches, the optimized results still provide high prediction accuracies for the validation data (92.2% accuracy) and the TDP-43 data (81.2% accuracy), serving as validation for the effectiveness of the descriptors used in ECLiPSE. It is important to note that the developed descriptors based on mostly point clouds are useful to quantitatively describe the data, and are capable of identifying the geometric differences between the distinct groups in the data, even when no information on the classes is provided to the model.

For the tau aggregates data, the situation is more complicated, as the within-class variance is much larger and between-class variance much smaller. Consequently, the prediction power of the most optimal unsupervised approach for this data is much lower than for the other two data sets (66.6% accuracy). This is still better than complete random predictions, especially because three of the four classes are much higher than the expected 25% for random prediction. This means that the unsupervised clustering approach recognizes the natural grouping in the data to some extent, but the model would not lead to accurate conclusions if it was used for quantification of the tau degradation experiment.

#### Supervised classification

Just as for the unsupervised classification, many different supervised classification methods were used in the process and evaluated with respect to each other. The different available methods in ECLiPSE are Binary/Multiclass Adaptive Boosting[13], K-nearest neighbors[14], Adaptive Logistic Regression[15], Random Forest[16], and Random Undersampling Boosting[17] from the MATLAB Statistics and Machine Learning Toolbox. Furthermore, two other methods, Logistic Regression [18] and Partial Least Squares classification[19], were implemented using the available functions of the PLS Toolbox (Eigenvector Inc.). It is worth noting that the linear discriminant analysis classification method[20] was not included in ECLiPSE because it assumes that the variables are normally distributed, which can oftentimes not be guaranteed, and it is sensitive to outliers. However, it is available in the MATLAB Statistics and Machine Learning toolbox and is an excellent classification method when all its assumptions are met.

In this work, the average prediction accuracy is used as a benchmark for which method performs better as it can be directly compared between the different methods if it is applied on the same training and validation data. For each of the data sets presented in the paper, 1,000 models were trained on a subset of the entire data, and then the 100 best models, defined as the models with the highest average prediction accuracy, were automatically selected and combined into an aggregated model to further improve performance. Training multiple classification models also provides an uncertainty on its performance, which can subsequently be considered when selecting the best classification model(s). It can for example be beneficial to select a model that is slightly less accurate in average prediction accuracy but provides a more robust prediction (i.e., lower standard deviation among the models). The results of the supervised methods that were considered for the three different data sets are shown in Supplementary Figure 3. A general conclusion of the results is that the difference in classification performance for the different methods is not large, which is an indication that the developed descriptors to quantify the data are well representative of the clusters and the differences between them, again validating their use.

Before discussing the analysis strategy and results in more detail for each data set, it is worth noting that a hyperparameter optimization study was performed for each of the methods to determine the best settings, and results are summarized in Supplementary Table 2.

#### Validation data

In this data set, each of the 1,000 models was trained on a subset of the data, comprising 553 randomly selected clusters for each category, which amounts to 50% of the smallest class size. While the default setting in ECLiPSE allocates 75% of the smallest class size for training data, we adjusted this proportion to 50% for the validation data, given the ease of distinguishing between various groups in this data set. This modification was made to enhance the models' generalizability. By training models on fewer data points, the variability between the training and validation data sets in the 1,000 distinct models increases, ultimately leading to more robust models. The validation data set was created by also randomly selecting 553 clusters for each type. Within each of the classification models, the training and validation data sets were autoscaled independently from each other, a necessary step in the classification to remove differences in scale between the different variables. To allow a direct comparison between non-variable selected and variable selected data, the models were trained and validated on the same subsets of the data.

The classification results (Supplementary Figure 3A) obtained on the validation data set are largely similar for each of the different methods used, with a small advantage in prediction power for the Random Forest method on the non-variable selected data (97.1 ± 0.1% accuracy). For this data, it is worth noting that the classification on the variable selected data performs similarly to the non-variable selected data because the complexity of the data is relatively low, and the different groups could easily be separated using just a limited number of components in the PCA/PLS space.

#### Tau aggregates data

Contrary to the validation data, the ground truth of this data set is not known; therefore, a small part of the acquired data was manually annotated into one of the four distinct aggregate classes: linear fibrils, branched fibrils, pre-NFTs, or NFTs. Many different strategies were considered when optimizing the classification results for this data set because of its complexity and results were evaluated with respect to each other. For example, besides equal class sizes for training and validation data sets, unbalanced training data sets (i.e., data sets in which the number of data points for each class is different) were also considered, but led to slightly inferior results. Finally, most optimal results with smallest prediction uncertainties were obtained by including 75% of the smallest class size (corresponding to 722 data points) of each cluster type. The validation data set then consisted of 241 clusters for each type (25% of the smallest class size). Again, both training and validation data sets used in each of the 1,000 classification models were autoscaled independently.

The results for all classification methods applied on this data are shown in Supplementary Figure 3B. It is clear from this figure that, due to the complexity of the data, performing variable selection before classification provided better results (except for Partial Least Squares classification). Moreover, it also improved the robustness of the models as the uncertainties are smaller. Additionally, because the uncertainties on the results obtained with Logistic Regression on the variable selected data (89.8 ± 0.4% accuracy) are small, this method was used for prediction rather than Multiclass Adaptive Boosting (90.2 ± 0.6% accuracy) or Random Forest (90.3 ± 0.5% accuracy) as the confidence on the results of the individual classes is higher. The 100 best models of this method were used to predict the aggregate type of each of the clusters imaged in the degradation study. Two autoscaling options were considered before applying the classification models on this data (i.e., considering each day individually, or combining the data). Both approaches provided the same results, which indicates once again that the developed methodology and classification models are robust and can be generalized.

#### TDP-43 data

The last data set presented in this work is the TDP-43 data set. The ground truth of this data set is known, so no manual annotation had to be performed. However, compared to the validation data, the structures in this data are not entirely distinct and thus the data is more complicated. For this reason, the classification model training was also performed by using 75% of the smallest class, which corresponds to 517 clusters. The validation data set was constructed using 173 clusters of each type (25% of the smallest data class). Training and validation data sets were autoscaled as previously, and the different methods were evaluated using the same subsets as well.

The full results for this data set are shown in Supplementary Figure 3C, which show once again that the classification method used only has a small influence on the final classification results, and that the variable selected data always led to better classification results (except for Partial Least Squares classification, which provided the highest prediction accuracy).

### Supplementary Note 3 – Comparison to state-of-the-art classification

To establish the validity of ECLiPSE, it is compared to a state-of-the-art classification technique called Automated Structures Analysis Program (ASAP)[21]. ASAP requires that localizations are rendered by a Point Spread Function (PSF) with a given width to produce images, which are also binarized by applying a threshold. Descriptors are then calculated based of these rendered and binarized images. We compare our results to those from ASAP using the validation data (Supplementary Figure 4) and the tau aggregation data (Figure 2A).

#### Influence of the user-provided parameters

##### Influence on descriptors

For ASAP, the user must provide the PSF width with which the localizations are rendered into an image, as well as a threshold that transforms the rendered image into a binary image. These additional input parameters have an influence on the resulting images, as indicated in Supplementary Figure 5. The PSF width has an influence on the amount of detail that is captured in the image, where a larger PSF width will result in the loss of small spatial details and provide smoother images. The threshold determines what cut-off is used to establish which intensity of the pixels is considered 'information' or not. It is therefore dependent on the number and density of localizations (as this determines the intensity of the rendered image pixels). These parameters should ideally be optimized for each cluster separately, or, considering that spatial scales and localization density is uniformly distributed within an experiment, they should be optimized for each experiment. This approach can potentially be easily optimized for simple and uniform structures like the nuclear pore complex. However, for complex data that spans a large range of length and density scales, like the tau data in this manuscript, the optimization needs to be done on a structure-by-structure basis, which becomes prohibitively difficult and time consuming. An example of the effects of the user-provided parameters is shown in Supplementary Figure 5, for a mitochondrion and lysosome. The figure illustrates the smoother images obtained with the larger rendering PSF width, as well as the significant impact of the different binary image threshold on the selected pixels for the mitochondrion cluster. On the other hand, the binary images of the lysosome cluster remain largely unaffected for the same threshold values. Hence, the optimization is highly dependent on the specific structure and a one-size-fits-all set of parameters cannot be applied universally to all data sets.

##### Influence on classification results

To further show the influence of the user-provided parameters for ASAP, the classification results on a subset of the lysosomes and mitochondria clusters (500 total clusters for each type) are shown in Supplementary Figure 6 (using the default ASAP Discriminant Analysis classification method). These results show that the influence is varying, and no clear trends can be noticed. In the case of the 10 nm PSF width, the threshold varies by ~3% over the investigated range. However, for the 20 nm PSF width, the results are largely independent on the threshold used. The results differ by ~3% when comparing different PSF widths as well. It is therefore important to spend a substantial amount of time exploring and optimizing a range of different parameters to maximize results, which is highly time consuming. An extended study on the influence of these two parameters was also performed for all different classification methods available in ASAP (i.e., Discriminant Analysis, K-nearest Neighbors, Naïve Bayes, and Classification Trees), and the results are presented in Supplementary Table 3. There is no clear trend in how the rendering PSF width influences the classification results, as results get better with an increasing rendering PSF width for the Discriminant Analysis method, but worsen for the K-nearest Neighbor and Naïve Bayes methods, and remain relatively stable for the Classification Trees method. Note also that the best results are obtained using a different threshold, even within the same classification method, with again no clear trend observable.

#### Classification comparison

##### Validation data

The classification accuracy of both ASAP and ECLiPSE was evaluated on the Lysosome vs Mitochondria data set. For both methods, default settings were used (Logistic Regression and no variable selection for ECLiPSE, and 10 nm rendering PSF width and $1.5 \times 10^5$ threshold for ASAP). The difference in average classification accuracy between this data set was 6.1% in favor of ECLiPSE (96.5% accuracy vs 90.4% accuracy), but, when comparing to the best ASAP results presented in Supplementary Table 3 (i.e., after extensive optimization), the average ECLiPSE classification accuracy is still 2.6% better (96.5% accuracy vs 93.9% accuracy).

When considering the full validation data set, similar results are achieved (Figure 1G): ECLiPSE performs 3.4% better on average and handles the correct classification of the challenging Mitochondria clusters much better, with a difference of 13.5%.

##### Tau aggregation data

The tau aggregates data set, which includes 200 clusters of each of the four different aggregate types (Linear Fibrils, Branched Fibrils, Pre-NFTs, and NFTs) was also used to compare ECLiPSE to ASAP, as this is a more challenging dataset for classification than the validation data. The differences between results obtained with ECLiPSE and ASAP ranges from a minimum of 3.8% difference in accuracy for the Linear Fibrils vs NFTs data set to a maximum of 17.7% difference in accuracy for the Linear Fibrils vs Branched Fibrils vs Pre-NFTs data set (Supplementary Table 4), all in favor of ECLiPSE.

### Supplemental Materials and Methods

#### Sample preparation and image acquisition

##### Aggregates of tau protein

***Fixing and immunostaining.*** Stable human embryonic kidney-derived QBI-293 cells (Clone 4.1[22]) expressing full length human tau T40 (2N4R) carrying the P301L mutation with a GFP tag were grown in Dulbecco's Modified Eagle Medium (DMEM) supplemented with 10% tetracycline-screened fetal bovine serum (FBS), 1% pyruvate (10 mM), 1% penicillin-streptomycin and L-glutamine (20 mM), 5 µg $mL^{-1}$ blasticidin, 200 µg $mL^{-1}$ Zeozin and maintained in an incubator at 37 °C with 5% $CO_2$. Clone 4.1 was continuously maintained in media containing 100 ng $mL^{-1}$ Doxycycline (Dox+), or Dox was removed from the culture media for several days to perform experiments (Day 1 -Dox, Day 2 -Dox, etc.) and then fixed. Cells were then incubated with stabilizing buffer (MTSB: 50 mM PIPES, 5 mM EGTA, 5 mM $MgSO_4 . 7H_2O$, 90 mM KOH in distilled water, at pH 7) for 3 min and then methanol (ice cold) was added for 3 min. After that, cells were washed with MTSB twice, followed by blocking for 1 h using 4% (wt/vol) BSA in phosphate buffer saline (PBS). They were then immunostained with GFP VHH nanobody, recombinant binding protein (#GT-250, Chromotek) conjugated with AlexaFluor 647 in 4% (wt/vol) BSA and 0.2% Triton X-100 (vol/vol, Thermo Fischer Scientific) in PBS.

***STORM imaging.*** Tau aggregates data were imaged on the Oxford Nanoimager microscope (ONI, Oxford, UK) equipped with a 100 × oil immersion objective (NA 1.45), 405-, 488-, 561- and 640-nm lasers, 498-551- and 576-620-nm band-pass filters in channel 1, 666-705-839 nm band-pass filters in channel 2, and an 840 Hamamatsu Flash 4 V3 sCMOS camera. The STORM imaging buffer used contained 50 mM Tris, 10 mM NaCl, 0.5 mg $mL^{-1}$ glucose oxidase (Sigma, G2133), 40 µg $mL^{-1}$ catalase (Roche Applied Science, 106810), 10% (wt/vol) glucose, and 30 mM Ciseamine (stock, 77 mg $mL^{-1}$ of 360 mM HCl), at pH 7.5. Images were collected with a 15 ms exposure time for 50,000 frames with constant laser power. The localizations were generated and drift corrected using the Nanoimager operating and analysis software (ONI).

***Cluster segmentation.*** Tau aggregate localizations were Voronoi-segmented and clustered based on a maximum Voronoi area of 410 $nm^2$ and a minimum of 5 localizations using a custom-made MATLAB code (https://github.com/melikelakadamyali/StormAnalysisSoftware). Clusters with less than 500 localizations were additionally filtered and removed from the analysis to investigate only the larger tau aggregate species (corresponding to larger linear fibrils, branched fibrils, pre-NFTs, and NFTs).

##### Nuclear Pore Complex

The preparation and acquisition of the nuclear pore complex (NPC) data has been described previously[23]. Below is a summary.

***Fixing and immunostaining***. U-2 OS genome-edited Nup96-mEGPF cells (clone 195,300174, CLS Cell Lines Service) were grown at 37 °C with 5% $CO_2$ in DMEM, to which MEM NEAA, GlutaMAX and 10% PBS was added. The cells were then fixed in PBS containing 4% (vol/vol) paraformaldehyde for 25 min and blocked for 1 h using 3% BSA and 0.2% triton X-100 in PBS. The cells were then immunostained with GFP VHH nanobody, recombinant binding protein (#GT-250, Chromotek) conjugated with AlexaFluor 647, and then washing for four times using washing buffer (0.2% blocking buffer and 0.05% Triton X-100 in PBS) for 10 min.

***STORM imaging.*** NPCs were imaged on the Oxford Nanoimager microscope (ONI, Oxford, UK) as described before. The STORM imaging buffer was as described above. Images were collected with a 10 ms exposure time for 40,000 frames with constant laser power. The localizations were generated and drift corrected by Nanoimager operating and analysis software (ONI).

***Cluster segmentation.*** NPC localizations were Voronoi-segmented and clustered based on a maximum of 821 $nm^2$ Voronoi area and a minimum of 75 localizations using a custom-made MATLAB code (https://github.com/melikelakadamyali/StormAnalysisSoftware). Clusters with an area smaller than 0.011 $\mu m^2$ and greater than 0.023 $\mu m^2$ were discarded as non-specific background clusters.

#### Microtubules

***Fixing and immunostaining***. BSC-1 cells (ATCC) were permeabilized for 10-30 s in buffer containing 80 mM PIPES-KOH pH 7.1, 1 mM EGTA, 1 mM $MgCl_2$, 0.5% Triton X-100 (vol/vol), 10% glycerol (vol/vol), followed by fixation in PBS containing 3% paraformaldehyde and 0.1% glutaraldehyde at 37 °C for 10 min. The fixed cells were washed twice with PBS before incubation with 0.1% (wt/vol) sodium borohydride for 7 min at 25 °C and washed again three times with PBS. Cells were incubated with blocking buffer (PBS containing 10% (vol/vol) donkey serum, 0.2% (vol/vol) Triton X-100, and 0.05 mg $mL^{-1}$ sonicated salmon sperm single stranded DNA (Stratagene, La Jolla, California) in PBS) before incubation with mouse anti-acetylated α-tubulin antibody (#48389, Abcam) at 1:100 dilution in blocking buffer for 1 h at 25 °C or overnight at 4 °C. The excess antibody was removed by three washes in 1× Wash Buffer (Massive Photonics, Germany) before incubating with docking-strand-conjugated secondary anti-mouse antibody (Docking strand 1, Massive Photonics) at 1:100 dilution in Antibody Incubation Buffer (Massive Photonics) for 1 h at 25 °C. The excess secondary antibody was removed by three washes with 1× Wash Buffer and twice with PBS.

***DNA-PAINT imaging.*** Acetylated microtubules were imaged on the Oxford Nanoimager microscope (ONI, Oxford, UK) as described before. ATTO-655 conjugated Imager 1 strands (Massive Photonics) were added to imaging chamber at 0.5 nM concentration in Imaging Buffer (Massive Photonics). Images were collected at HiLo illumination angle with continuous imaging mode at 100 ms exposure time for 10,000 frames at 30 °C. The localizations were generated and drift corrected using the Nanoimager operating and analysis software (ONI), and localizations beyond 30 nm precision and outside the 10-150 nm $\sigma_{X/Y}$ range were removed from downstream analysis.

***Cluster segmentation.*** Microtubule localizations were Voronoi-segmented and clustered based on a maximum Voronoi area of 5,475.6 $nm^2$ and a minimum of 5 localizations using a custom-made MATLAB code (https://github.com/melikelakadamyali/StormAnalysisSoftware). Clusters with an area smaller than 0.041 $\mu m^2$ were discarded as non-specific background clusters. Sections of microtubules were selected as regions of interest only from well-separated single microtubules.

#### Lysosomes

***Fixing and immunostaining.*** HeLa cells were fixed in PBS containing 4% paraformaldehyde warmed to 37 °C for 20 minutes at 25 °C. Fixed cells were washed three times with PBS, followed by permeabilization in 0.1% (vol/vol) Saponin in PBS. Cells were then incubated in blocking buffer (PBS containing 10% (vol/vol) donkey serum, 0.1% (vol/vol) Saponin, and 0.05 mg $mL^{-1}$ sonicated salmon sperm single stranded DNA (Stratagene) for one hour at 25 °C before incubation with mouse anti-LAMP2 (sc18822, Santa Cruz) at 1:100 dilution in blocking buffer for 1 h at 25 °C. The excess antibody was removed by three washes in 1×

Wash Buffer (Massive Photonics) before incubating with docking-strand-conjugated secondary anti-mouse antibody (Docking strand 1, Massive Photonics) at 1:100 dilution in Antibody Incubation Buffer (Massive Photonics) for 1 h at 25 °C. The excess secondary antibody was removed by three washes with 1× Wash Buffer and twice with PBS.

***DNA-PAINT imaging.*** Lysosomes were imaged on the Oxford Nanoimager microscope (ONI, Oxford, UK) as described before. Cy3B or ATTO-655 conjugated Imager 1 strands (Massive Photonics) were added to imaging chamber at 0.5 nM concentration in Imaging Buffer (Massive Photonics). Images were collected at HiLo illumination angle with continuous imaging mode at 100 ms exposure for 25,000 frames. Localizations were generated and drift corrected using the Nanoimager operating and analysis software (ONI).

***Cluster segmentation.*** Lysosome localizations were Voronoi-segmented and clustered based on a maximum Voronoi area ranging from 342-684 $nm^2$ and a minimum of 25 localizations using a custom-made MATLAB code (https://github.com/melikelakadamyali/StormAnalysisSoftware). Lysosome clusters with and area less than 0.123 $\mu m^2$ and greater than 0.479 $\mu m^2$ were discarded as non-specific background clusters or clustering artefacts.

#### Mitochondria

***Fixing and immunostaining.*** Cos7 cells were fixed in PBS containing 4% paraformaldehyde warmed to 37 °C for 20 minutes at 25 °C. Fixed cells were washed three times with PBS, followed by permeabilization in 0.2% (vol/vol) Triton X-100 in PBS. Cells were then incubated in blocking buffer (PBS containing 10% (vol/vol) donkey serum, 0.2% (vol/vol) Triton X-100, and 0.05 mg $mL^{-1}$ sonicated salmon sperm single stranded DNA (Stratagene) for one hour at 25 °C before incubation with rabbit anti-Tom20 (11802-1-AP, Proteintech) at 1:100 dilution in blocking buffer for 1 h at 25 °C or overnight at 4 °C. The excess antibody was removed by three washes in 1× Wash Buffer (Massive Photonics) before incubating with docking-strand-conjugated secondary anti-rabbit antibody (Docking strand P1[24]) at 1:25 or 1:100 dilution in Antibody Incubation Buffer (Massive Photonics) for 1 h at 25 °C. The excess secondary antibody was removed by three washes with 1× Wash Buffer and twice with PBS.

***DNA-PAINT imaging.*** Mitochondria were imaged on the Oxford Nanoimager microscope (ONI, Oxford, UK) as described before. Cy3 conjugated Imager P1 strands (IDT) were added to imaging chamber at 0.5 nM concentration in Imaging Buffer (Massive Photonics). Images were collected at HiLo illumination angle with continuous imaging mode at 10 ms exposure for 50,000 frames. Localizations were generated and drift corrected using the Nanoimager operating and analysis software (ONI).

***Cluster segmentation.*** Mitochondria localizations were Voronoi-segmented and clustered based on a maximum Voronoi area ranging from 178-2738 $nm^2$ and a minimum of 25 localizations using a custom-made MATLAB code (https://github.com/melikelakadamyali/StormAnalysisSoftware). Mitochondria clusters with an area less than 0.274 $\mu m^2$ and greater than 1.03 $\mu m^2$ were discarded as non-specific background clusters or clustering artefacts.

#### TDP-43

***Brain seeds.*** Sarkosyl-insoluble TDP-43 protein from the frozen frontal cortex of FTLD-TDP patients was prepared as described previously[25]. Human postmortem brains were obtained from the University of Pennsylvania, CNDR Brain Bank[26]. All necessary written informed consent forms were obtained from the patients or their next of kin in accordance with University of Pennsylvania Institutional Review Board guidelines and confirmed at the time of death.

Brain-derived TDP-43 extracts from FTLD-TDP cases used in the present study have been previously characterized by Porta *et al.*, 2021 (Cases #4 and #12)[27].

***Cellular TDP-43 aggregation assay***. iGFP-NLSm cells (clone #6.B7) were plated on coated poly-D-lysine chambered-coverglass Nunc Labtek II (16,000 cells/well) and transduced after 24h with 0.5 µg brain-derived TDP-43 extracts (100-300 pg TDP-43/well)[25,27]. Briefly, brain extracts were sonicated and diluted with dPBS and mixed with single-use tubes of BioPORTERTM as a protein delivery reagent (BP509696, Genlantis) as previously described[25,27]. Protein–Bioporter complexes were added to the cells and incubated for 4 h. Cells were placed back on fresh medium in presence of doxycycline (1.0 µg/ml) and cultured for 3 additional days.

***Immunocytochemistry.*** To remove cytoplasmic soluble proteins and visualize the formation of phosphorylated TDP-43 aggregates, transduced iGFP-NLSm were fixed in 4% paraformaldehyde (PFA) containing 1% Triton X-100 for 15 min at room temperature. Briefly, after blocking cells were incubated with the mAb phospho-specific p409-410 antibody (1:5,000, 80007-1-RR, Proteintech) overnight at 4°C. After three washes with dPBS cells were incubated with anti-rabbit Alexa 647 conjugated 1:100 in 4% BSA in PBS 1× for 1 h at room temperature in the dark. Stained cells were rinsed with PBS containing 2% BSA and 0.5% Tx-100 and keep in PBS at 4 °C.

***Imaging.*** Strain-specific TDP-43 aggregates were imaged on the Oxford Nanoimager Microscope (ONI, Oxford, UK) as described before, with an imaging buffer as described above as well. Each field of view was imaged with a 15 ms exposure time for 25,000 frames at constant laser power and at a HiLo angle of illumination. The localizations were generated and drift corrected using the Nanoimager operating and analysis software (ONI).

***Cluster segmentation.*** The phosphorylated TDP-43 localizations were Voronoi-segmented and clustered based on a maximum Voronoi area of 684 nm$^2$ (Strain A) and 1.37 µm$^2$ (Strain B) and a minimum of 10 localizations using a custom-made MATLAB code (https://github.com/melikelakadamyali/StormAnalysisSoftware). Additionally, clusters with an area smaller than 0.05 µm$^2$ were discarded.

#### Data sets

##### Validation data

The validation data set was constructed by combining 1190 Tau, 1660 NPCs, 1328 Microtubules, 1106 Lysosome, and 1181 Mitochondria structures. The tau structures were randomly selected from the manually annotated *branched fibrils* class of the tau aggregates data.

##### Tau aggregates data

This data set consists of data that originate from three biological replicates for each of the imaged days in the degradation process. The total number of data is: 11263 clusters for +Dox control (29 cells), 7461 clusters for Day 1 -Dox (27 cells), 7517 clusters for Day 2 -Dox (30 cells), 3583 clusters for Day 3 -Dox (27 cells), 1981 clusters for Day 4 -Dox (29 cells), 1865 clusters for Day 5 -Dox (27 cells), and 288 clusters for Day 10 -Dox (28 cells).

To construct the data set for classification model training, a subset of the data (representing ~30% of the data from these days) from the +Dox control, Day 1 -Dox, and Day 2 -Dox imaging days was manually annotated. To avoid bias in this process, the data was randomized, and the person annotating was blinded from the origins of the data. After annotation, 963 linear fibrils, 5218 branched fibrils, 1098 pre-NFTs, and 1350 NFTs were identified.

#### TDP-43 data

The TDP-43 data was constructed by combining the data from the two different strains of TDP-43 (strain A and strain B). The total number of clusters was 900 (15 fields of view) and 694 (19 fields of view), respectively.

### Supplementary Figures

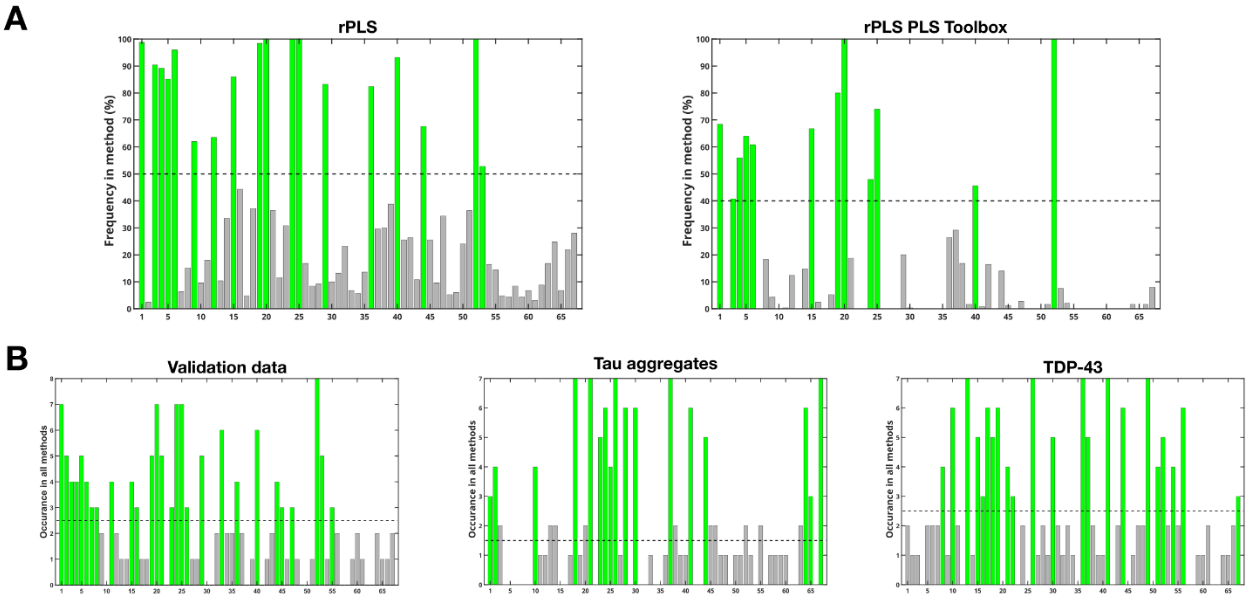

**Supplementary Figure 1 Variable selection results.** (A) Results of the two recursive Partial Least Squares routines with different implementations applied on the validation data, showing the stricter implementation in the PLS Toolbox; (B) The number of times a variable was selected using the different routines (left panel: validation data – 28 total variables selected; middle panel: Tau aggregate data – 27 total variables selected; right panel: TDP-43 data – 22 total variables selected). The dashed line in each plot represents the cutoff threshold for which variables are important. Corresponding variables can be found in Supplementary Table 1.

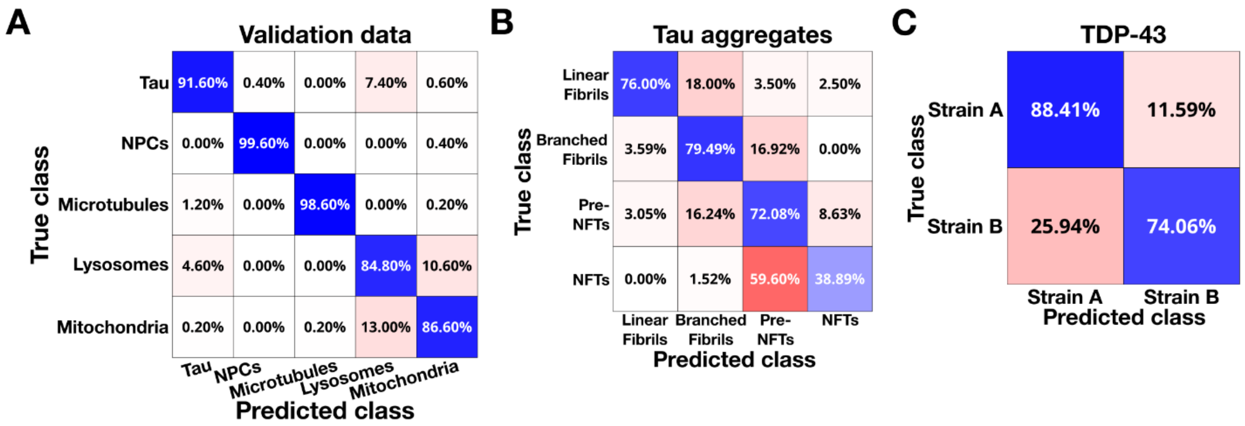

**Supplementary Figure 2 Results obtained by unsupervised classification (agglomerative hierarchical clustering) after optimization.** (A) Validation data without variable selection: Ward's method – PCA transformation (18 principal components) – Euclidean distance (92.2% accuracy); (B) Tau aggregates data with variable selection: Ward's method – PCA transformation (4 principal components) – Euclidean distance (66.6% accuracy); (C) TDP-43 data with variable selection: Ward's method – PCA transformation (1 principal component) – Mahalanobis distance (81.2% accuracy).

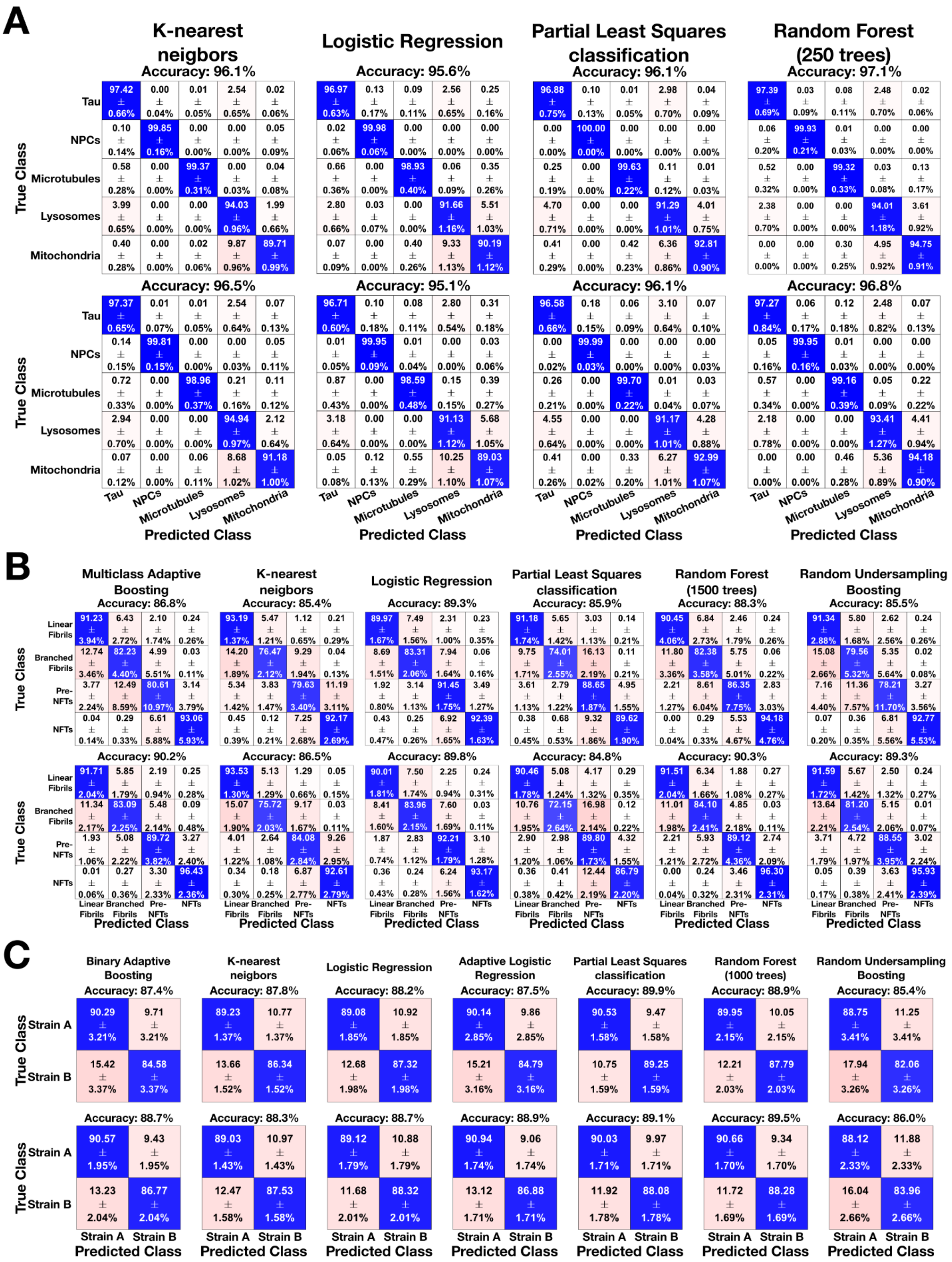
A
K-nearest neigbors
Accuracy: 96.1%
Logistic Regression
Accuracy: 95.6%
Partial Least Squares classification
Accuracy: 96.1%
Random Forest (250 trees)
Accuracy: 97.1%
True Class
Tau
NPCs
Microtubules
Lysosomes
Mitochondria
Accuracy: 96.5%
Accuracy: 95.1%
Accuracy: 96.1%
Accuracy: 96.8%
Predicted Class
B
Multiclass Adaptive Boosting
Accuracy: 86.8%
K-nearest neigbors
Accuracy: 85.4%
Logistic Regression
Accuracy: 89.3%
Partial Least Squares classification
Accuracy: 85.9%
Random Forest (1500 trees)
Accuracy: 88.3%
Random Undersampling Boosting
Accuracy: 85.5%
Linear Fibrils
Branched Fibrils
Pre-NFTs
NFTs
Accuracy: 90.2%
Accuracy: 86.5%
Accuracy: 89.8%
Accuracy: 84.8%
Accuracy: 90.3%
Accuracy: 89.3%
C
Binary Adaptive Boosting
Accuracy: 87.4%
K-nearest neigbors
Accuracy: 87.8%
Logistic Regression
Accuracy: 88.2%
Adaptive Logistic Regression
Accuracy: 87.5%
Partial Least Squares classification
Accuracy: 89.9%
Random Forest (1000 trees)
Accuracy: 88.9%
Random Undersampling Boosting
Accuracy: 85.4%
Strain A
Strain B
Accuracy: 88.7%
Accuracy: 88.3%
Accuracy: 88.7%
Accuracy: 88.9%
Accuracy: 89.1%
Accuracy: 89.5%
Accuracy: 86.0%

**Supplementary Figure 3 Results obtained by supervised classification after optimization**. (A) the validation data without (top) and with (bottom) variable selection. The best result is obtained with Random Forest (97.1 ± 0.1% accuracy) on the non-variable selected data; (B) the tau aggregates data without (top) and with (bottom) variable selection. The best result is obtained using Random Forest (90.3 ± 0.5% accuracy) on the variable selected data; (C) the TDP-43 data without (top) and with (bottom) variable selection. The best results are obtained using Partial Least Squares classification (89.9 ± 0.6% accuracy) on the non-variable selected data. Note that the Logistic Regression results are also shown in Supplementary Figure 7.

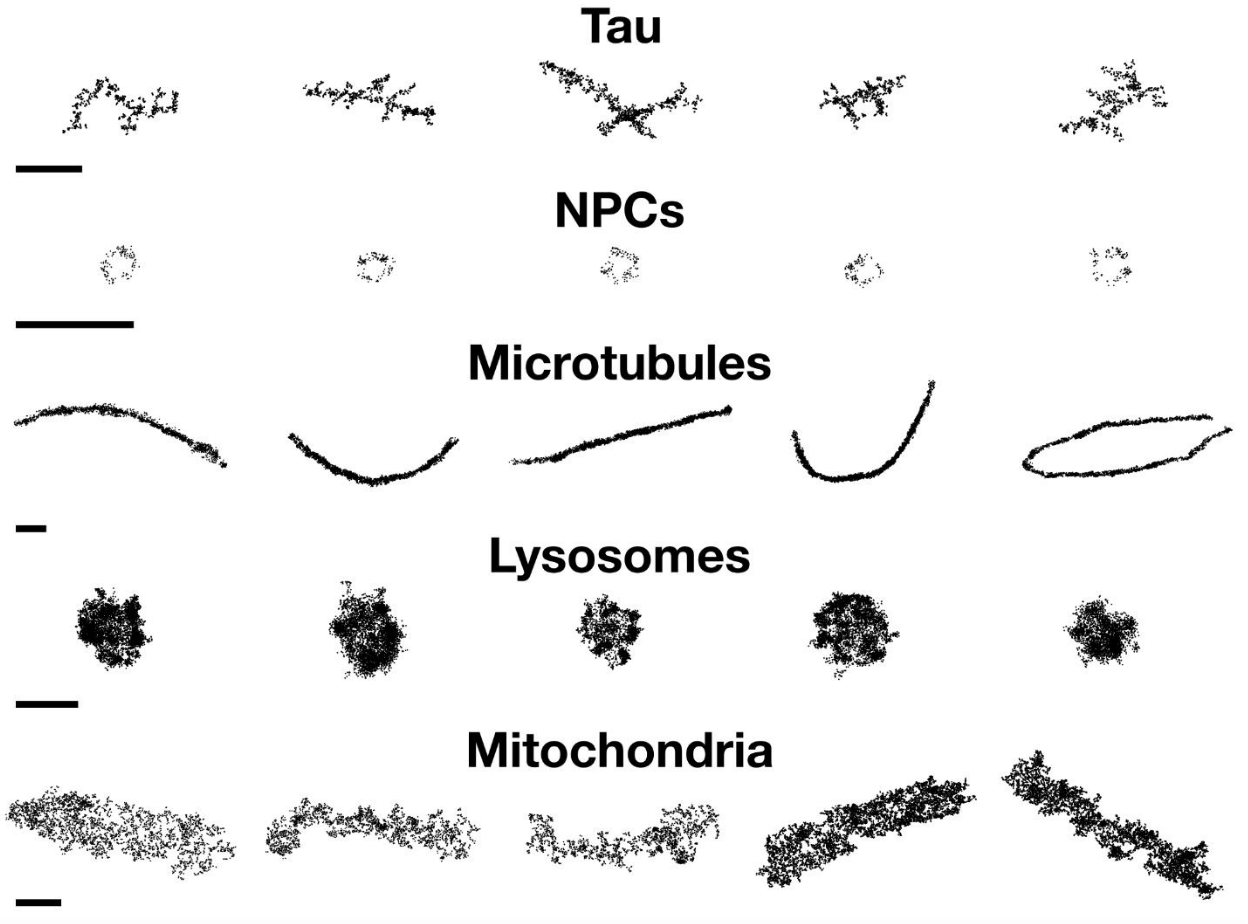

**Supplementary Figure 4 Example clusters of the five different biological structures included in the validation data:** Tau clusters (1190 clusters), Nuclear Pore Complex clusters (1660 NPCs), Microtubule clusters (1328 Microtubules), Lysosome clusters (1106 Lysosomes), and Mitochondrion clusters (1181 Mitochondria). Scale bar is 500 nm.

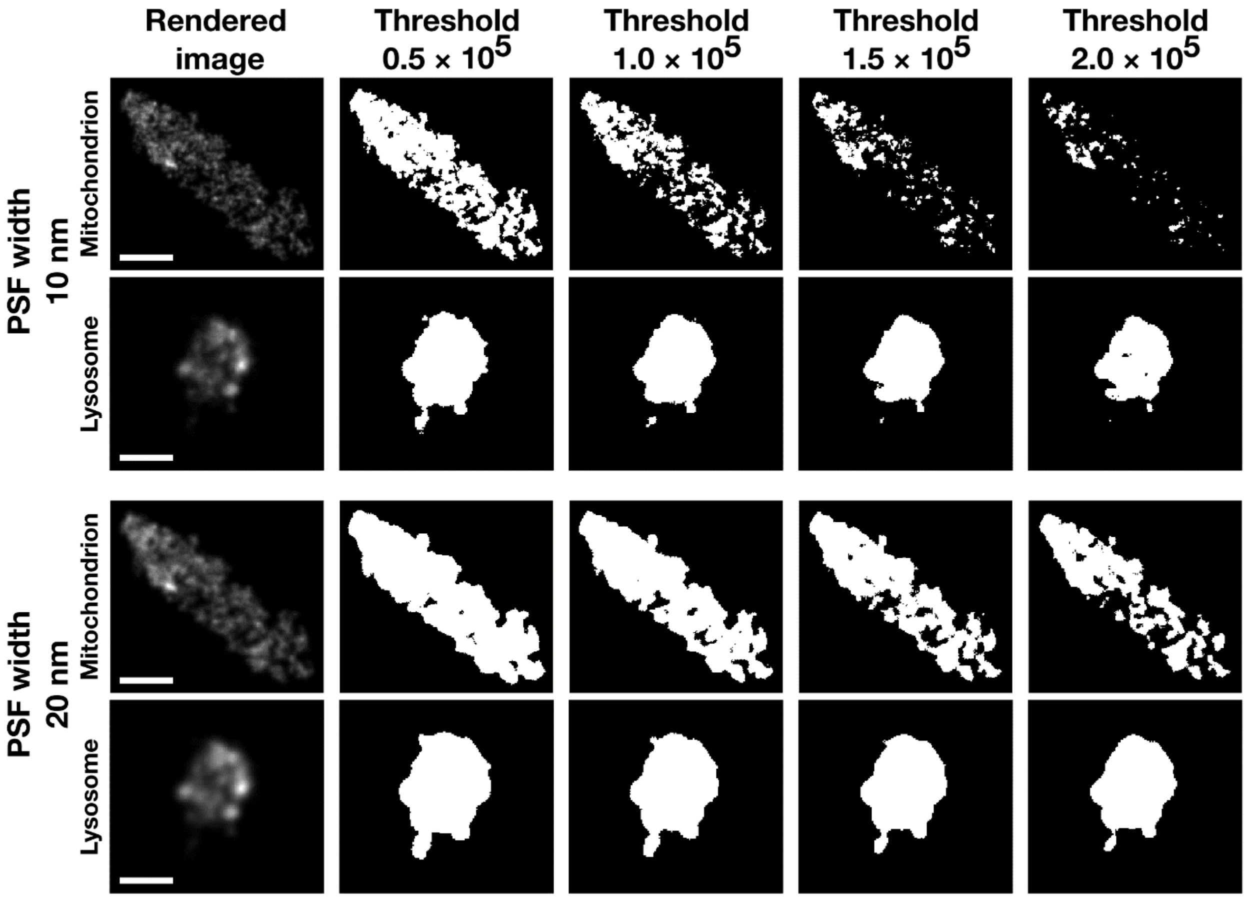

**Supplementary Figure 5 The effect of the rendering PSF width and binary image threshold on the ASAP images, applied to a mitochondrion and a lysosome.** A larger rendering PSF width 'smooths' out the image and structure, and fine details are lost, whereas an increased threshold leads to removal of signal of the mitochondrion, but does not have a large influence for the lysosome. Scale bar is 500 nm.

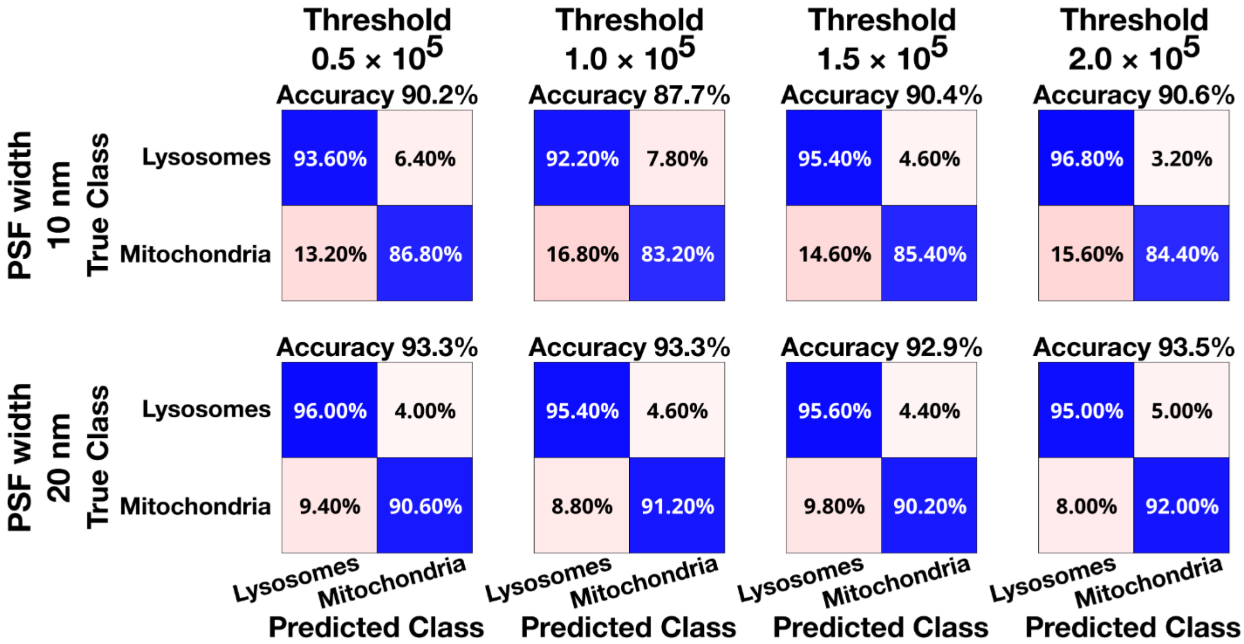

**Supplementary Figure 6 Results obtained using ASAP (default Discriminant Analysis classification method) on the Lysosomes – Mitochondria data.** Both the rendering PSF width and the binary image threshold have an influence on the classification results.

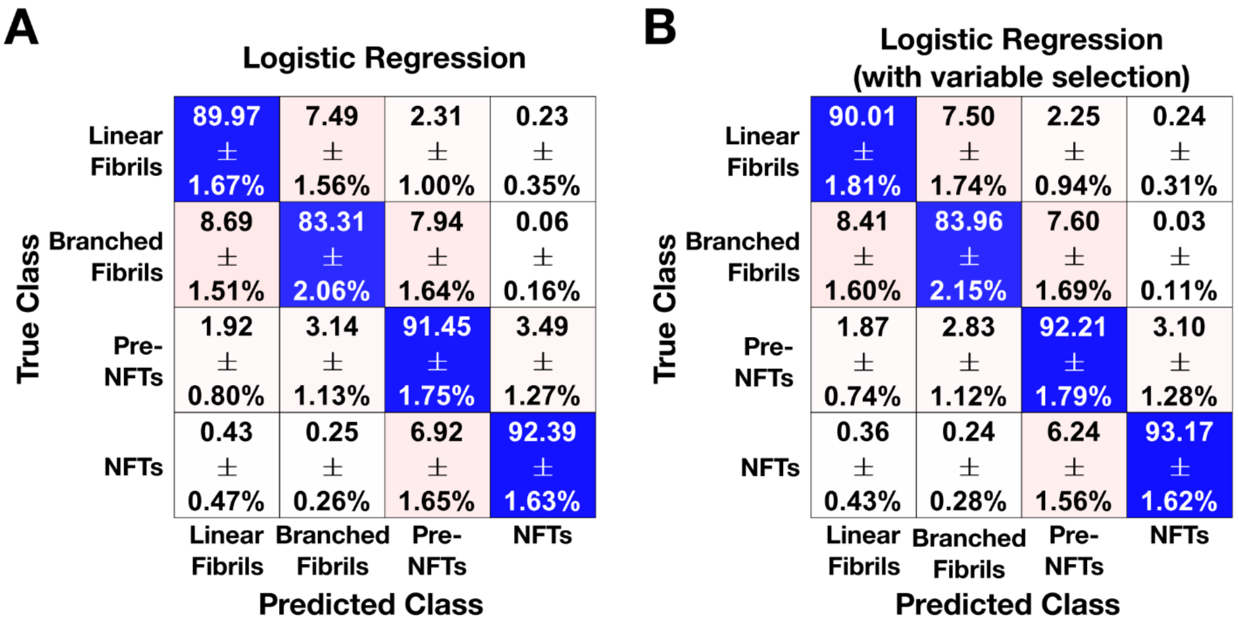

**Supplementary Figure 7 Comparison between the results obtained using Logistic Regression on the tau aggregates data,** (A) without (88.3 ± 0.4% accuracy) and (B) with variable selection (89.8 ± 0.4% accuracy). Note that these results are also included in Supplementary Figure 3 for comprehensiveness.

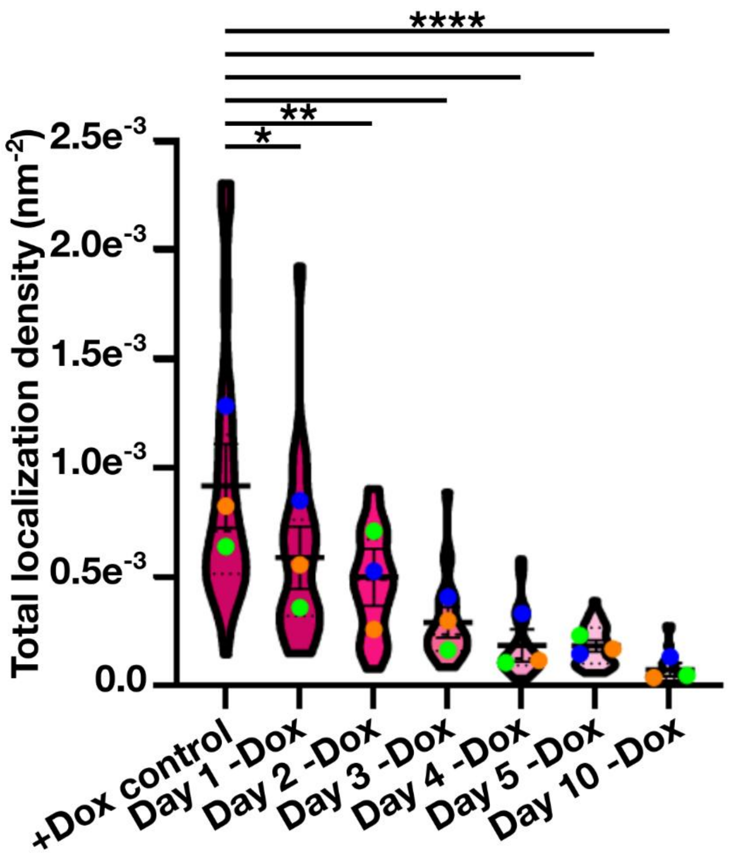

**Supplementary Figure 8 Quantification of the total localization density of tau proteins in cells in control cells (+Dox) and various days following Dox removal**. The different colored circles represent three biological replicates, full lines indicate the median, dotted lines indicate the 25$^{th}$ and 75$^{th}$ percentile. +Dox control: n = 29 cells, Day 1 -Dox: n = 27 cells, Day 2 -Dox: n = 30 cells, Day 3 -Dox: n = 27 cells, Day 4 -Dox: n = 29 cells, Day 5 -Dox: n = 27 cells, Day 10 -Dox: n = 28 cells. *: $p < 0.05$, **: $p < 0.001$, ****: $p < 0.0001$.

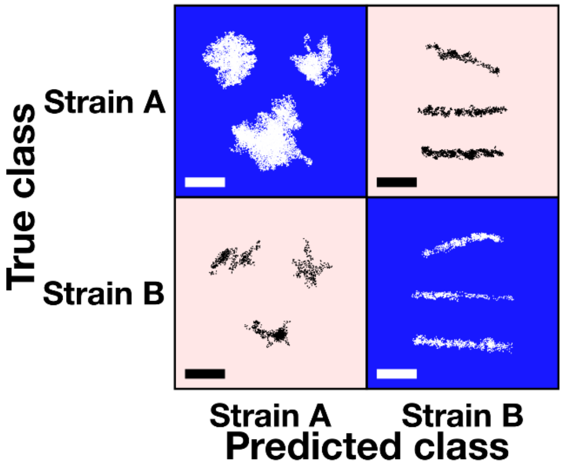

**Supplementary Figure 9 Representative images of correctly and wrongly classified TDP-43 clusters for the two available strains (Strain A and strain B).** The misclassified clusters have the morphological character of the other strain showing the mixed character of each strain. Scale bar is 500 nm.

#### Supplementary Tables

**Supplementary Table 1 Details on the type and implementation of the different descriptors used to quantitatively describe the SMLM point clouds.**

| **30 Geometric descriptors** | | | |
|---|---|---|---|
| ***#*** | ***Name*** | ***Description*** | ***Implementation*** |
| 1 | Number of localizations | The total number of localizations inside the data | Point cloud |
| 2 | Major axis | The major axis of the data (largest range) | Point cloud |
| 3 | Minor axis | The minor axis of the data (smallest range) | Point cloud |
| 4 | Area | The total area of the data | Alphashape of point cloud |
| 5 | Filled area | The total area of the data in which holes (within boundaries of the data) are filled up | Polygon of point cloud |
| 6 | Convex area | The area of the convex hull of the data<br>(Convex hull: the smallest set of points on the boundary of the data delimiting a region containing all boundary points) | Polygon of point cloud |
| 7 | Radius of Gyration | The radial distance of a circle with the same moment of inertia as the data | Point cloud |
| 8 | Euler Number | The total number of objects minus the total number of holes in the data | Polygon of point cloud |
| 9 | Eigenvalues ratio | The ratio between the x and y eigenvalues | Point cloud |
| 10 | Eigenentropy | Information entropy that represents a measure for order/disorder of two-dimensional points within the covariance ellipsoid. | Point cloud |
| 11 | Density | The density of the data | Point cloud + Alphashape of point cloud |
| 12 | Area ratio | The ratio between the area of the data including holes and the filled area | Alphashape of point cloud + Polygon of point cloud |
| 13 | Aspect ratio | The ratio between the major and minor axis | Point cloud |
| 14 | Form ratio | The ratio between the area and the square of the major axis | Alphashape of point cloud + Point cloud |

Supplementary Table 1 Continued

| # | Name | Description | Implementation |
|---|---|---|---|
| 15 | Rectangularity | The ratio between the area and the minimum bounding box (also known as Extent) | Alphashape of point cloud + Point cloud |
| 16 | Circularity | A measure for how round the data is | Polygon of point cloud |
| 17 | Convex circularity | A measure for how circular the convex hull of the data is | Polygon of point cloud |
| 18 | Convexity | A measure for how convex the data is with respect to the convex hull | Polygon of point cloud |
| 19 | Solidity | A measure for the proportion of the data filling the convex hull of the data | Alphashape of point cloud + Polygon of point cloud |
| 20 | Equivalent diameter | The diameter of the circle with the same area as the data | Polygon of point cloud |
| 21 | Fiber length | A measure that represents the length of the fiber if the data was a fiber | Alphashape of point cloud |
| 22 | Fiber width | A measure that represents the width of the fiber if the data was | Alphashape of point cloud |
| 23 | Curl | A measure for how curled the data is | Alphashape of point cloud + Point cloud |
| 24 | Major axis normalized to area | The major axis normalized by the total area of the data | Alphashape of point cloud + Point cloud |
| 25 | Minor axis normalized to area | The minor axis normalized by the total area of the data | Alphashape of point cloud + Point cloud |
| 26 | Ellipse major axis | The length of the major axis of the ellipse with the same normalized second central moment as the data | Alphashape of point cloud |
| 27 | Ellipse minor axis | The length of the minor axis of the ellipse with the same normalized second central moment as the data | Alphashape of point cloud |
| 28 | Ellipse aspect ratio | The aspect ratio between the ellipse major and minor axis | Alphashape of point cloud |
| 29 | Ellipse ratio | A measure for how elliptical the data is | Alphashape of point cloud + Polygon of point cloud |
| 30 | Ellipse eccentricity | The ratio of the distance between the foci of the ellipse and the major axis of that ellipse. | Alphashape of point cloud |

Supplementary Table 1 continued

| # | Name | Description | Implementation |
|---|---|---|---|
| | | **7 Boundary descriptors** | |
| ***#*** | ***Name*** | ***Description*** | ***Implementation*** |
| 31 | Full perimeter | The perimeter of the data, including the perimeter where holes are inside the data | Alphashape of point cloud |
| 32 | Outer perimeter | The outer perimeter of the data (as in accordance with the 'Filled area' descriptor) | Polygon of point cloud |
| 33 | Convex perimeter | The perimeter of the convex hull of the data | Polygon of point cloud |
| 34 | Elastic energy | The elastic energy of the boundary points of the data | Point cloud |
| 35 | Bending energy | The bending energy of the boundary points of the data | Point cloud |
| 36 | Mean curvature | The mean curvature of the boundary points of the data | Point cloud |
| 37 | Bending energy normalized to area | The bending normalized by the total area of the data | Alphashape of point cloud + Point cloud |
| | | **8 Skeleton descriptors** | |
| ***#*** | ***Name*** | ***Description*** | ***Implementation*** |
| 38 | Median skeleton width | The median width of the data | High-definition Alphashape binarization of point cloud |
| 39 | Total skeleton length | The total length of the skeleton of the data | High-definition Alphashape binarization of point cloud |
| 40 | Number of skeleton segment intersections | The total number of intersections (i.e., points where segments touch) in the skeleton of the data | High-definition Alphashape binarization of point cloud |
| 41 | Mean length of skeleton segments | The mean length of the segments of the skeleton of the data (a substitute measure how 'branched' the data is) | High-definition Alphashape binarization of point cloud |
| 42 | Mean orientation of skeleton segments | The mean orientation of each skeleton segment with respect to the major axis of the data | High-definition Alphashape binarization of point cloud |
| 43 | Mean tortuosity of skeleton segments | The mean tortuosity (i.e., ratio between the length and Euclidean distance between start and end point) of each skeleton segment | High-definition Alphashape binarization of point cloud |

Supplementary Table 1 continued

| # | Name | Description | Implementation |
|---|---|---|---|
| 44 | Total skeleton length normalized to area | The total skeleton length normalized by the area | High-definition Alphashape binarization of point cloud + Alphashape of point cloud |
| 45 | Number of skeleton segment intersections normalized to area | The number of skeleton segment intersections normalized to the area | High-definition Alphashape binarization of point cloud + Alphashape of point cloud |
| | | **9 Texture descriptors** | |
| ***#*** | ***Name*** | ***Description*** | ***Implementation*** |
| 46 | Contrast | A measure of intensity contrast between data points and their neighbors | High-definition 3D histogram of the data |
| 47 | Correlation | A statistical measure of correlation between data points and their neighbors | High-definition 3D histogram of the data |
| 48 | Energy | A measure for the uniformity of the data | High-definition 3D histogram of the data |
| 49 | Entropy | A statistical measure of randomness that characterizes texture of the data | High-definition 3D histogram of the data |
| 50 | Kurtosis | A statistical measure for whether the data is peaked or flat with respect to a normal distribution | High-definition 3D histogram of the data |
| 51 | Mean intensity | The mean intensity of the data | High-definition 3D histogram of the data |
| 52 | Mean non-zero intensity | The mean intensity of all non-zero contributions in the data | High-definition 3D histogram of the data |
| 53 | Root Mean Square Roughness | A statistical measure for the roughness of the data | High-definition 3D histogram of the data |
| 54 | Skewness | A measure of lack of symmetry of the data | High-definition 3D histogram of the data |
| | | **7 HuMoment descriptors** | |
| ***#*** | ***Name*** | ***Description*** | ***Implementation*** |
| 55 | First HuMoment | First Invariant with respect to scale, rotation and translation.<br>Analogous to the moment of inertia, and this moment is reflection symmetric. | High-definition Alphashape binarization of point cloud |

Supplementary Table 1 continued

| # | Name | Description | Implementation |
|---|---|---|---|
| 56 | Second HuMoment | Second Invariant with respect to scale, rotation and translation (reflection symmetric). | High-definition Alphashape binarization of point cloud |
| 57 | Third HuMoment | Third Invariant with respect to scale, rotation and translation (reflection symmetric). | High-definition Alphashape binarization of point cloud |
| 58 | Fourth HuMoment | Fourth Invariant with respect to scale, rotation and translation (reflection symmetric). | High-definition Alphashape binarization of point cloud |
| 59 | Fifth HuMoment | Fifth Invariant with respect to scale, rotation and translation (reflection symmetric). | High-definition Alphashape binarization of point cloud |
| 60 | Sixth HuMoment | Sixth Invariant with respect to scale, rotation and translation (reflection symmetric). | High-definition Alphashape binarization of point cloud |
| 61 | Seventh HuMoment | Sevent Invariant with respect to scale, rotation and translation (reflection anti-symmetric). | High-definition Alphashape binarization of point cloud |

**6 Fractal descriptors**

| # | Name | Description | Implementation |
|---|---|---|---|
| 62 | Local Minkowski Bouligand dimension | Fractal dimension of the data that measures how the complexity of details changes with scale (also known as box counting) | High-definition Alphashape binarization of point cloud |
| 63 | Minkowski sausage | The dimension representing the fractal resulting from replacing each side of a square by broken lines | High-definition Alphashape binarization of point cloud |
| 64 | Hausdorff dimension | A measure for the roughing using fractals | High-definition Alphashape binarization of point cloud |
| 65 | Boundary Hausdorff dimension | Hausdorff dimension of the boundary | High-definition Alphashape binarization of point cloud |
| 66 | Skeleton Minkowski sausage | Minkowski sausage of the skeleton | High-definition Alphashape binarization of point cloud |
| 67 | Skeleton Hausdorff dimension | Hausdorff dimension of the skeleton | High-definition Alphashape binarization of point cloud |

**Supplementary Table 2 Optimized hyperparameters for each classification method per data set.**

| | | Validation data | Tau aggregates data | TDP-43 data |
|---|---|---|---|---|
| Binary/multiclass Adaptive Boosting | Learning rate | - | 0.1 | 0.1 |
| | Learning cycles | - | 1,000 | 1,000 |
| | Splits | - | 10 | 10 |
| K-nearest neighbors | Neighbors | 5 | 15 | 15 |
| | Distance metric | Standardized Euclidean | cosine | Cosine |
| Logistic Regression | Compression | PLS | PLS | PLS |
| | Components | 5 | 15 | 5 |
| | Cross validation | Venetian blinds 5 groups 1 blind size | Venetian blinds 5 groups 1 blind size | Venetian blinds 5 groups 1 blind size |
| Adaptive Logistic Regression | Learning rate | - | - | 0.1 |
| | Learning cycles | - | - | 1,000 |
| | Splits | - | - | 10 |
| Partial Least Squares classification | Algorithm | SIMPLS | SIMPLS | SIMPLS |
| | Orthogonalization | No | No | No |
| | Components | Automatically optimized per round | Automatically optimized per round | Automatically optimized per round |
| Random Forest | Trees | 250 | 1,500 | 1,000 |
| Random Undersampling Boosting | Learning rate | - | 0.1 | 0.1 |
| | Learning cycles | - | 1,000 | 1,000 |
| | Splits | - | 10 | 10 |

**Supplementary Table 3 Influence of the rendering PSF width and binary image threshold on the classification results aimed at separating Lysosomes and Mitochondria.** Four different values for the binary image threshold were used ($0.5 \times 10^5$, $1.0 \times 10^5$, $1.5 \times 10^5$, and $2.0 \times 10^5$) and the best classification accuracy obtained is reported in this table. For reference, an accuracy of 96.5% is obtained using the ECLiPSE descriptors and default classification method (Logistic Regression, no variable selection).

| | Rendering PSF width | | | |
|---|---|---|---|---|
| | 10 nm (%) | 15 nm (%) | 20 nm (%) | 25 nm (%) |
| Discriminant Analysis | $90.4^3$ | $92.5^1$ | $93.5^4$ | $93.9^2$ |
| K-nearest Neighbors | $90.0^3$ | $89.3^4$ | $88.8^4$ | $85.9^1$ |
| Naïve Bayes | $81.9^1$ | $80.7^1$ | $80.1^1$ | $79.2^1$ |
| Classification trees | $92.2^4$ | $92.5^4$ | $90.9^1$ | $91.2^4$ |

Best result obtained with $^1$: $0.5 \times 10^5$ threshold; $^2$: $1.0 \times 10^5$ threshold; $^3$: $1.5 \times 10^5$ threshold; $^4$: $2.0 \times 10^5$ threshold

**Supplementary Table 4 Comparison between the results obtained on the tau aggregate data using the default method and settings in ECLiPSE and default ASAP method and settings.** The smallest difference in results observed is for the LFs vs NFTs data set, whereas the largest difference in results is observed in the LFs vs BFs vs Pre-NFTs data set.

| | ECLiPSE$^A$ (%) | ASAP$^B$ (%) | Difference (%) |
|---|---|---|---|
| LFs vs BFs | 99.0 | 83.0 | +16.0 |
| LFs vs NFTs | 100.0 | 96.2 | + 3.8 |
| LFs vs BFs vs Pre-NFTs | 95.2 | 77.5 | +17.7 |
| BFs vs Pre-NFTs vs NFTs | 93.2 | 83.1 | +10.1 |
| LFs vs BFs vs Pre-NFTs vs NFTs | 92.9 | 80.6 | +12.3 |

$^A$: Logistic Regression classification, no variable selection

$^B$: PSF width: 10nm, $1.5 \times 10^5$ Threshold, Discriminant classification

LFs: Linear Fibrils; BFs: Branched Fibrils; Pre-NFTs: Precursor Neurofibrillary Tangles; NFTs: Neurofibrillary Tangles

#### Supplementary Videos

**Supplementary Video 1:** Principal Component Analysis (3 PCs are shown) applied to the validation data shows that the developed descriptors can discriminate well between five different classes. The left video shows PCA applied to the non-variable selected data (using all 67 descriptors; 62.9% explained variance) in which classes are well-separated but show a large within-class variance. The video on the right shows PCA applied to the variable selected data (using only 28 descriptors; 76.5% explained variance) demonstrating that the within-class variance is heavily reduced without compromising the separation between the different classes.
